# Functional MRI Signatures of Autonomic Physiology in Aging

**DOI:** 10.1101/2025.04.05.647057

**Authors:** Jiawen Fan, Meher R. Juttukonda, Sarah E. Goodale, Shiyu Wang, Csaba Orbán, Divya Varadarajan, Jonathan R. Polimeni, Catie Chang, David H. Salat, Jingyuan E. Chen

## Abstract

While traditionally regarded as “noise”, blood-oxygenation-level-dependent (BOLD) fMRI fluctuations coupled to systemic physiology—such as heart rate and respiratory changes—also hold valuable information about brain vascular properties and autonomic function. In this study, we leverage these physiological BOLD signals to characterize age-related changes in brain physiology. Using a large dataset from the Lifespan Human Connectome Project Aging study, we investigated how the spatiotemporal BOLD-fMRI signatures of autonomic physiology, specifically heart rate and respiratory variation, change with advancing age. Our findings reveal that aging is associated with globally slower respiratory fMRI responses, alongside faster cardiac fMRI responses and enhanced brain-cardiac signal coupling. Moreover, we show that the impact of age on physiological fMRI signals exhibits a notable turning point after age 60, suggesting a critical role of declining vascular health and autonomic function in aging. The potential impact of age-related changes in brain structure, tissue perfusion, and in-scan arousal states on the identified physiological fMRI patterns is also tested and discussed. Altogether, our results underscore significant age effects in the fMRI signatures of systemic physiology, emphasizing the pivotal role of altered vascular properties and autonomic function in aging. Methodologically, this study also demonstrates the utility of resting-state fMRI for extracting multi-parametric information about brain physiology, offering new biomarker opportunities that complement established functional connectivity metrics.

## Introduction

Hemodynamic changes that underlie the blood-oxygenation-level-dependent (BOLD) contrast in functional Magnetic Resonance Imaging (fMRI) are not only sensitive to regional neuronal activity but also influenced by systemic physiological processes regulated by the autonomic nervous system, including slow variations in respiratory depth, heart rate, and blood pressure [1–10]. Such BOLD fluctuations driven by various physiological processes are often considered as “noise” and removed from the fMRI time series during preprocessing.

While BOLD physiological dynamics may act as “noise” that confound the extraction of neuronal information from fMRI data, they also provide valuable insights into cerebral blood flow regulation, vascular anatomy and associated biomechanical properties, presenting an excellent opportunity to infer brain physiology in both health and disease. For instance, recent studies have proposed using respiratory “noise” in resting-state fMRI signals to characterize cerebrovascular reactivity (CVR)—the ability of cerebral blood vessels to dilate or constrict in response to vasoactive factors—and thereby inferring brain-wide vascular reserve [11,12]. Unlike conventional studies that employ hypercapnic CO2 inhalation [13–15], endogenous CO2 fluctuations driven by temporal variations in respiration can be leveraged to map CVR [11,12,16–18]. Beyond vascular reserve, the timing of respiratory responses—determined by the transit speed of CO2 from the lungs to brain tissues and by regional vasogenic responses, i.e., the arteriolar dilation triggered by elevated partial pressure of CO2 in arterial blood—also provides valuable information about arterial transit time and, ideally, regional delays in hemodynamic responses [19].

In addition to respiratory “noise”, there is also rich information regarding brain-heart interactions and the vascular mechanical properties embedded in the cardiac “noise” in the form of widespread fMRI fluctuations following a change in heart rate [3,5]. Distinct from active vasogenic responses to CO2 changes, these cardiac fMRI responses are linked to beat-to-beat variations in the blood “piped” from the heart through the brain, and thus primarily track passive vasodilation due to arterial compliance caused by blood pressure waves. As such, the timing and amplitude of cardiac fMRI responses may serve as markers of arterial stiffness, offering insights into the biomechanical properties of the arteries.

The goal of this study is to leverage the rich, multi-parametric information embedded in physiological fMRI “noise” to gain insights into age-dependent changes in the autonomic regulation and vascular physiology. Previous research has shown that aging could lead to changes in the tortuosity of penetrating arterioles [20–22] as well as a reduction in cerebral capillary and arteriole density in older brains [23–29], which could cause slower blood transit times from large arteries to capillaries [30] and reduced vascular reactivity to systemic physiological stimuli such as changes in CO₂ levels and other vasodilators [31–34]. Additionally, age-related mechanism could alter biomechanical properties of the blood vessels, resulting in increased arterial stiffness [35]. This stiffening is attributed to the accumulation of collagen and degradation of elastin fibers within the vessels [36], which in turn reduce cerebral blood flow and cerebrovascular reserve, impair deep white matter perfusion, and compromise vascular integrity [37–40]. These age-related changes in vascular health and autonomic physiology likely manifest in the timing and amplitudes of systemic physiological fMRI “noise”, suggesting that it may be possible to infer the degradation of vascular or autonomic function from dynamic fMRI data. Moreover, these physiological fMRI metrics offer advantages over conventional physiological imaging techniques—such as MR-based perfusion and angiography methods or transcranial Doppler—by providing superior spatiotemporal resolution, higher sensitivity, and whole-brain coverage. Surprisingly, existing fMRI-based aging studies often neglect the altered physiological components and predominantly focus on the neurogenic functional activity reflected in the fMRI time-series data [41,42].

Here, by analyzing a large, public cohort of resting-state fMRI datasets from the Lifespan Human Connectome Project Aging (HCP-A) study [43,44], we conduct a comprehensive evaluation of the impact of aging on fMRI signatures of autonomic physiology. By linking brain-wide fMRI dynamics with low-frequency peripheral respiratory and pulse measures, we show that aging induces significant changes in various physiological fMRI metrics, including globally slower fMRI responses driven by respiratory changes, faster fMRI responses following heart-rate variations, and an increased coupling between brain fMRI and cardiac signals. We further demonstrate that age-related alterations in these physiological fMRI metrics do not change linearly across the lifespan, with a distinct inflection in age dependence observed after age sixty. These physiological changes are interpreted in terms of age-related shifts in tissue perfusion and arterial compliance, and their connections to accompanying structural and arousal-state alterations are also examined and discussed. Together, our findings demonstrate and elucidate detailed changes in fMRI signatures of autonomic physiology in aging, highlighting altered vascular properties and autonomic function as significant features of late-life stages. Methodologically, our study also shows that resting-state physiological fMRI “noise” can be used to extract multi-parametric physiological information, providing a valuable complement to existing fMRI research that focuses primarily on brain neuronal function in clinical and neuroscientific contexts. A preliminary account of this study has been presented in abstract form [45].

## Results

### Subjects and Data

Resting-state fMRI data from a cohort of 400 adults from the HCP-A Lifespan 2.0 Release [43,44] were analyzed in this study to characterize changes in fMRI signatures of autonomic physiology in aging. This subset of subjects was chosen by visually inspecting the peripheral sensory recordings (pulse oximetry and respiratory bellows) to ensure the quality of extracted physiological metrics. Subjects were further divided into two age groups: *Aging I* (36–60 yrs, *N*=227, 125 females) and *Aging II* (60–90 yrs, *N*=173, 91 females). Age 60 was chosen as the cut-off threshold because it has been previously marked as a pivotal point in the altered rate of decline in brain structural and functional changes [46–50].

### Age effects on respiratory variation (RV)-coupled fMRI dynamics

We first examined how naturalistic respiratory responses—spatiotemporal dynamics of fMRI signals following an increase in respiratory variation—change with aging. RV was derived by computing time-windowed standard deviations of respiratory waveforms recorded by the bellows, tracking dynamic changes in respiratory volumes and depths [3]. A cross-correlation analysis was performed to characterize the temporal lags and couplings between brain-wide fMRI signals and RV.

Cross-correlations between the cerebral cortical fMRI signal, i.e., the mean across gray-matter voxels, and RV exhibited a bimodal pattern across all age groups—an increase in RV led to an initial overshoot, followed by a pronounced undershoot in the global fMRI signal (*Fig. 1a*). Note that here positive lag values correspond to the fMRI responses trailing the RV, and vice versa for the negative lag. A significant age effect was identified in the temporal lags associated with the trough of the cross-correlation results (i.e., corresponding to the undershoot in the respiratory fMRI response), with the elder group exhibiting more delayed respiratory responses (Aging I: 13.4 ± 3.9 s; Aging II: 16.3 ± 4.1 s, two-sample *t* test, *p* = 4.6 × 10^-12^). No significant between-group differences were observed in the correlation values associated with the trough (Aging I: –0.39 ± 0.17; Aging II: –0.39 ± 0.15, two-sample *t* test, *p* = 0.69), and no differences were observed in the peak of the cross-correlation plots including both the timing (Aging I: –0.06 ± 10.0 s; Aging II: –0.54 ± 8.39 s, two-sample *t* test, *p* = 0.61) and correlation value (Aging I: 0.24 ± 0.10; Aging II: 0.26 ± 0.11, two-sample *t* test, *p* = 0.074) (*Fig. 1a*).

**Figure 1.**
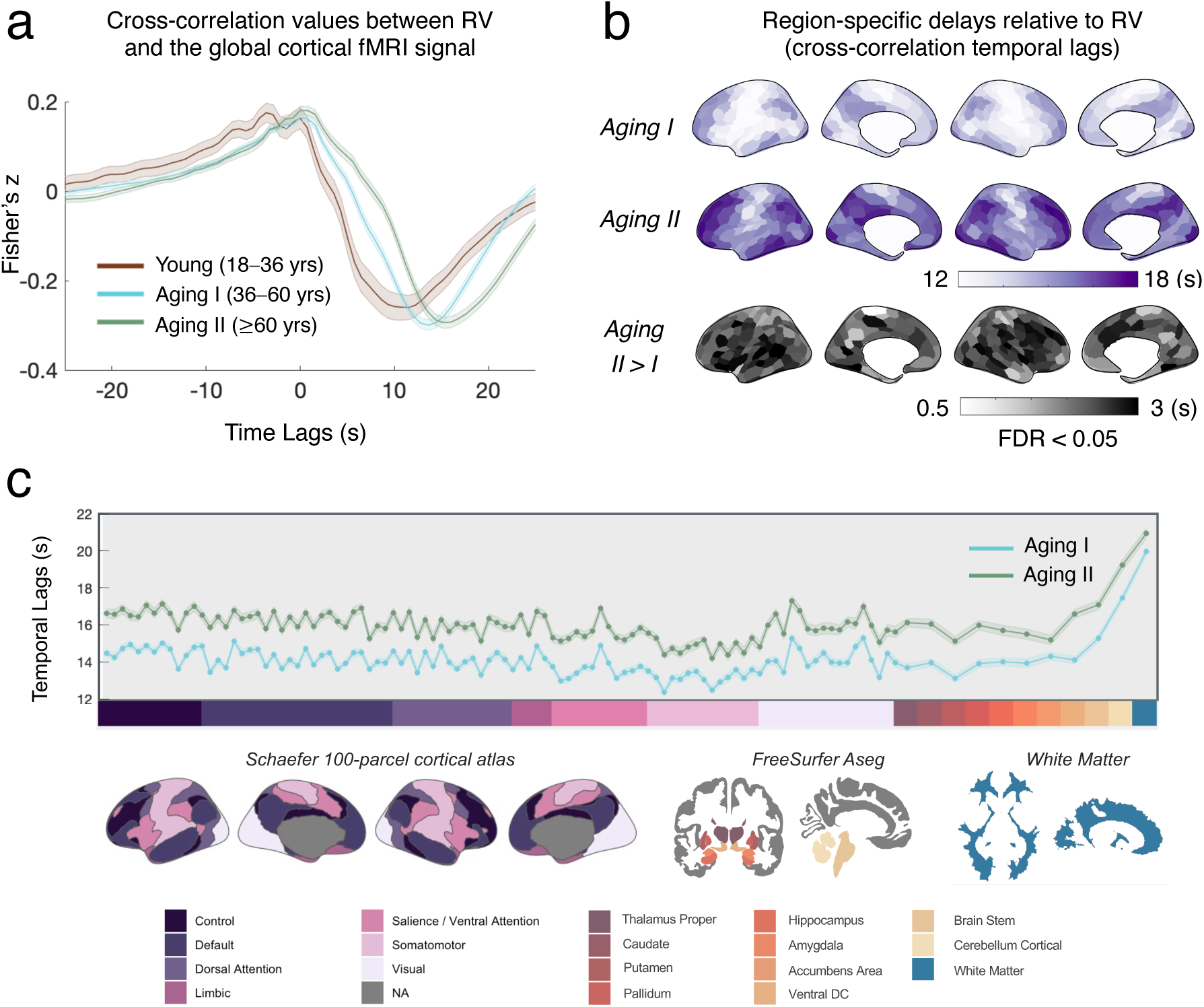
Age effects on the spatiotemporal patterns of RV-coupled fMRI dynamics. (a) The cross correlation between the global cortical fMRI signal and RV (positive lag values suggest that fMRI signals lag RV) for different age groups. The results of a cohort of 78 young healthy subjects from the HCP are also shown for comparison (“Young”, see *Supplementary Table S2* for detailed descriptions). Error bars indicate the standard errors of cross-correlation results across subjects within each age group. (b) Intra-cortical distributions of region-specific fMRI signal lag relative to RV (based on the Schaefer 300-parcel atlas). Regions exhibiting statistically significant between-group temporal lag differences were displayed at the bottom (“Aging II > I”, *p* < 0.05, FDR). (c) Tissue-type specific temporal lags relative to RV for each age group. Error bars indicate the standard errors of RV-fMRI temporal lags across subjects, and the shaded gray area highlights ROIs that exhibited statistically significant between-group differences (*p* < 0.05, FDR). ROI labels were shown at the bottom.

Consistent with our previous report on the HCP young-adult data (18–36 yrs) [51], the lifespan dataset also revealed spatially heterogenous temporal lags in the respiratory responses, quantified according to the trough of the cross-correlation results. Within the cortex, association cortex generally exhibited slower responses than the sensory cortex (*Fig. 1b*, Aging I and Aging II); and across the brain, the most delayed fMRI responses were identified in the cerebellum and white matter (*Fig. 1c*). As subjects aged, a brain-wide increase in the RV-fMRI temporal lags was observed, with the largest between-group lags reaching up to approximately 3 seconds (*Fig. 1b*, Aging II > I, & *1c*).

### Age effects on heart rate-coupled fMRI dynamics

We next investigated how cardiac responses—spatiotemporal dynamics of fMRI signals following an increase in heart rate—change with aging under resting state conditions. During the experiments, the aging group exhibited slightly slower heart rates and higher heart-rate variability after age 60 (*Fig. 2a,* and *Supplementary Fig. S1*), consistent with previous reports [52,53]. A time-varying waveform of the heart rate variation was derived by computing time-windowed averages of inter-heart-beat intervals (HBIs, i.e., the inverse of heart rate) recorded by pulse oximetry. Akin to the RV-fMRI analysis, cross-correlations were calculated to characterize the temporal lags and couplings between brain-wide fMRI signals and HBI.

**Figure 2.**
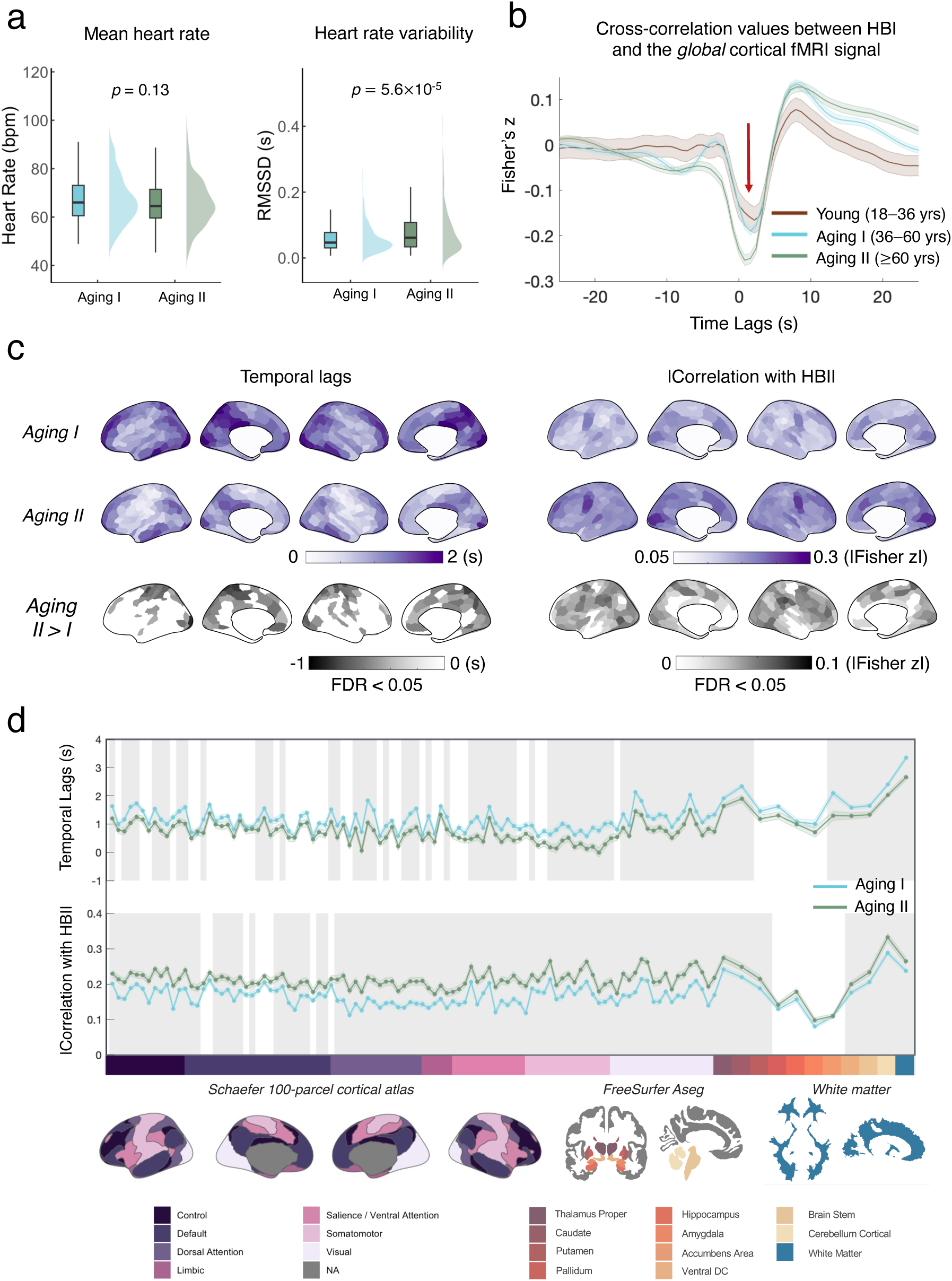
Age effects on the spatiotemporal patterns of heart rate-coupled fMRI dynamics. (a) Mean heart rate (left) and heart rate variability (right) of different age groups. (b) The cross correlation between the global cortical fMRI signal and HBI (positive lag values suggest that fMRI signals lag HBI) for different age groups. The results of a cohort of 78 young healthy subjects from the HCP are also shown for comparison (“Young”, see *Supplementary Table S2* for detailed descriptions). Error bars indicate the standard errors of cross-correlation results across subjects within each age group. (c) Intra-cortical distributions of region-specific fMRI signal lags (left) relative to and couplings (right) with HBI (based on the Schaefer 300-parcel atlas). Lag and peak correlation values were estimated based on the trough of cross-correlations (pointed by red arrow in (b)). Regions exhibiting statistically significant between-group temporal lag or correlation value differences were displayed at the bottom (“Aging II > I”, *p* < 0.05, FDR). (d) Tissue-type specific temporal lags (top) and cross-correlation peak correlation values (bottom) for each age group. Error bars indicate the standard errors of HBI-fMRI temporal lags and peak correlation values across subjects, and the shaded gray areas highlight ROIs that exhibited statistically significant between-group differences (*p* < 0.05, FDR). The labels and orders of ROIs are identical to Fig. 1c.

Age effects were also prominent in the cardiac responses. In the older cohort, we observed a faster cardiac response and increased coupling (i.e., stronger absolute correlation values) between global fMRI signals and the HBI when focusing on the trough of the cross-correlation (*Fig. 2b*, temporal lag: Aging I: 1.3 ± 1.4 s; Aging II: 0.9 ± 1.5 s, two-sample *t* test: *p* = 0.01; correlation value: Aging I: –0.22 ± 0.13; Aging II: –0.29 ± 0.15, two-sample *t* test: *p* = 2.0 × 10^-6^). No statistically significant between-group differences were identified for the later positive peak in the cross-correlation plots (temporal lag: Aging I: 8.9 ± 2.5 s; Aging II: 9.4 ± 3.0 s, two-sample *t* test, *p* = 0.041; peak correlation value: Aging I: 0.19 ± 0.14; Aging II: 0.20 ± 0.13, two-sample *t* test, *p* = 0.19).

Consistent with our previous report [51], cardiac responses in the lifespan cohort were also inhomogeneous across the brain (*Fig. 2c*, Aging I & II, & *2d*). The intra-cortical lag patterns of cardiac responses, quantified by the trough of the cross-correlation results, correlated positively with those of the respiratory responses (Spearman’s correlation *r* = 0.83, *p* < 10^-20^). For both age groups, the somatosensory and visual network consistently exhibited stronger HBI-fMRI coupling, while the limbic network (temporal pole and orbitofrontal cortex) displayed relatively low correlations with the heart rate signal.

Statistically significant reductions in the temporal lags and increases in the HBI-fMRI coupling with age were observed in most brain regions (*Fig. 2c, d*). However, the specific cortical patterns associated with the between-group difference were modestly correlated (Spearman’s correlation *r* = 0.20, *p* = 5.06×10^-4^), suggesting distinct mechanisms underlying age-related changes in the cardiac response timing and magnitudes.

### Linking age-related changes in physiological fMRI metrics to anatomical structure, tissue perfusion, and arousal states

Having identified significant age-related effects in both respiratory and cardiac fMRI responses, we next tested their associations with accompanying changes in brain structure, tissue perfusion, and arousal state dynamics during aging.

We first assessed whether the observed age-related effects in physiological fMRI metrics might partially stem from potential anatomical biases. For example, reduced percentage of gray matter within each fMRI voxel due to cortical atrophy [54,55], which may diminish the relative contributions of the neuronal component to the total variance of voxel-wise fMRI signals and thereby increases HBI-fMRI correlations. To test this hypothesis, we quantified the parcel-level gray-matter partial volumes using each subject’s high-resolution structural MRI data and compared the resulting spatial distributions with those of different physiological metrics. As expected, the regions showing the most significant reductions in gray matter percentage (*Fig. 3a*, “MGMF”) aligned with those reported to exhibit the most severe cortical thinning, including the frontoparietal cortex, limbic cortex, dorsal attention and somatomotor network [56–58]. However, the spatial pattern of reduced gray-matter partial volumes in aging was only modestly correlated with those of physiological metrics (*Fig. 3b*), suggesting that the partial volume effect had a limited influence on the observed age-related physiological patterns.

**Figure 3.**
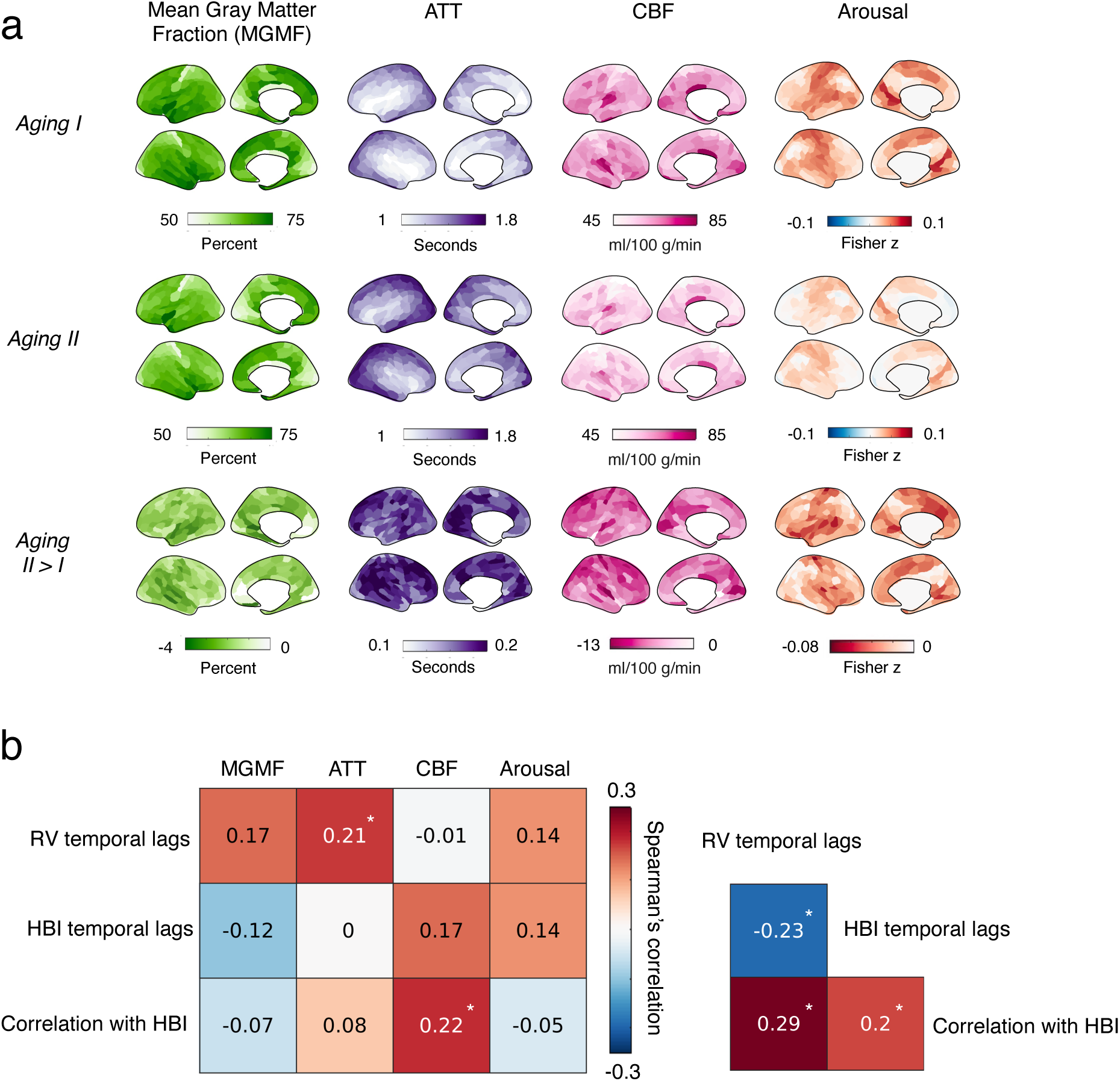
Spatial patterns of age effects: physiological fMRI metrics vs. structure, tissue perfusion, and arousal. (a) Group-level mean gray-matter fraction (MGMF), ATT, CBF, and eye camera-based arousal effects. “Aging I/II“: mean results within each age group; “Aging II > I“: spatial pattern of age effects. (b) Spearman’s spatial correlation of age effects between physiological fMRI metrics (Figs. 1b and 2c) and structure, blood perfusion and arousal effects (a). Pairwise correlation values were shown, with significant correlations indicated by an asterisk (*) following Bonferroni correction (*p* < 0.01).

We next examined the vascular physiological origins of the characterized respiratory and cardiac fMRI responses by linking their timing and amplitudes to two baseline physiological metrics—arterial transit time (ATT) and cerebral blood flow (CBF), estimated from arterial spin labeling (ASL) MRI data from the same subject cohort of the HCP-A project [30]. Since the timing of the BOLD responses to respiration is partially influenced by the speed of CO2 delivery, a positive coupling between RV-fMRI temporal lags and ATT was expected. Regarding cardiac BOLD responses, since previous studies have consistently reported reduced baseline CBF with increased arterial stiffness [37,59,60]—a key determinant of arterial compliance to the cardiac pressure wave—a positive association between CBF and cardiac responses is expected. Consistent with our predictions, the frontal-parietal regions showing the most significant between-group differences in respiratory responses also exhibited the most delayed ATT in aging (*Fig. 3*); and cortical regions with the most CBF reductions also exhibited faster cardiac responses and stronger HBI-fMRI coupling (*Fig. 3*). These findings corroborate the physiological basis of age-related changes in cardiac and respiratory fMRI responses identified in this study.

Finally, we tested the potential impact of in-scan vigilance levels on the coupling between systemic physiology and fMRI signals. This analysis was motivated by emerging evidence that arousal can intensify the apparent correlation between fMRI signals and physiological responses [61]. Thus, we assessed whether older subjects were drowsier during the experiments. Subjects’ vigilance levels during scans were inferred from eye camera data collected simultaneously during resting-state scans. For a subset of subjects (*N*=88) with high-quality eye camera data, we computed the instantaneous in-scan eye-closure frequency and correlated it with brain-wide fMRI signals to identify the fMRI signatures of vigilance. We considered this subset of subjects representative because the RV/HBI-fMRI coupling results derived from this subset were consistent with those from the full *N*=400 group (see *Supplementary Fig. S2a*). In both age groups, we observed modest effects of fluctuating arousal (see *Supplementary Fig. S2b*), likely due to the relatively short scan duration (∼6 minutes per fMRI scan). Contrary to our predictions, the older subjects exhibited a less pronounced arousal effect, evidenced by reduced correlations between fMRI signals and the eye-camera data (*Fig. 3*, “Arousal”). Additionally, the spatial alignment between age-related changes in arousal effects and HBI-fMRI correlation patterns was modest and statistically insignificant. Thus, differences in in-scan arousal levels are unlikely the primary drivers of the observed physiological fMRI changes in aging.

### Characterizing the “aging rate” of physiological fMRI metrics

As a final investigation, we examined whether age-related changes in various physiological fMRI metrics progress linearly as a function of age. This analysis was motivated by previous evidence that brain shrinkage, molecular profiling, and vascular aging change more quickly after age 60 [46–50].

Specifically, we tested whether the “aging rate”, defined as the linear dependence on age, of different physiological fMRI metrics altered after age 60 (i.e., Aging I vs. Aging II) using a linear regression model, and mapped brain regions exhibiting the most prominent between-group differences (see *Methods*). Apparent inflection in the linear dependence on age emerged for all physiological fMRI metrics after age 60 (*Fig. 4*). For the RV-fMRI temporal lag, increased aging rates were identified in midline structures (posterior cingulate cortex (PCC), anterior cingulate cortex (ACC), and precuneus), and at the temporal-frontal boundary (secondary somatosensory cortex (S2) and lateral sulcus). For the HBI-fMRI temporal lag, reduced aging rates were identified in the mid cingulate cortex (MCC), prefrontal cortex, and temporal lobe. Compared to the more focal patterns associated with the temporal lags, widespread cortical regions, except for a few visual and prefrontal patches, exhibited increased aging rates of HBI-fMRI correlation values.

**Figure 4.**
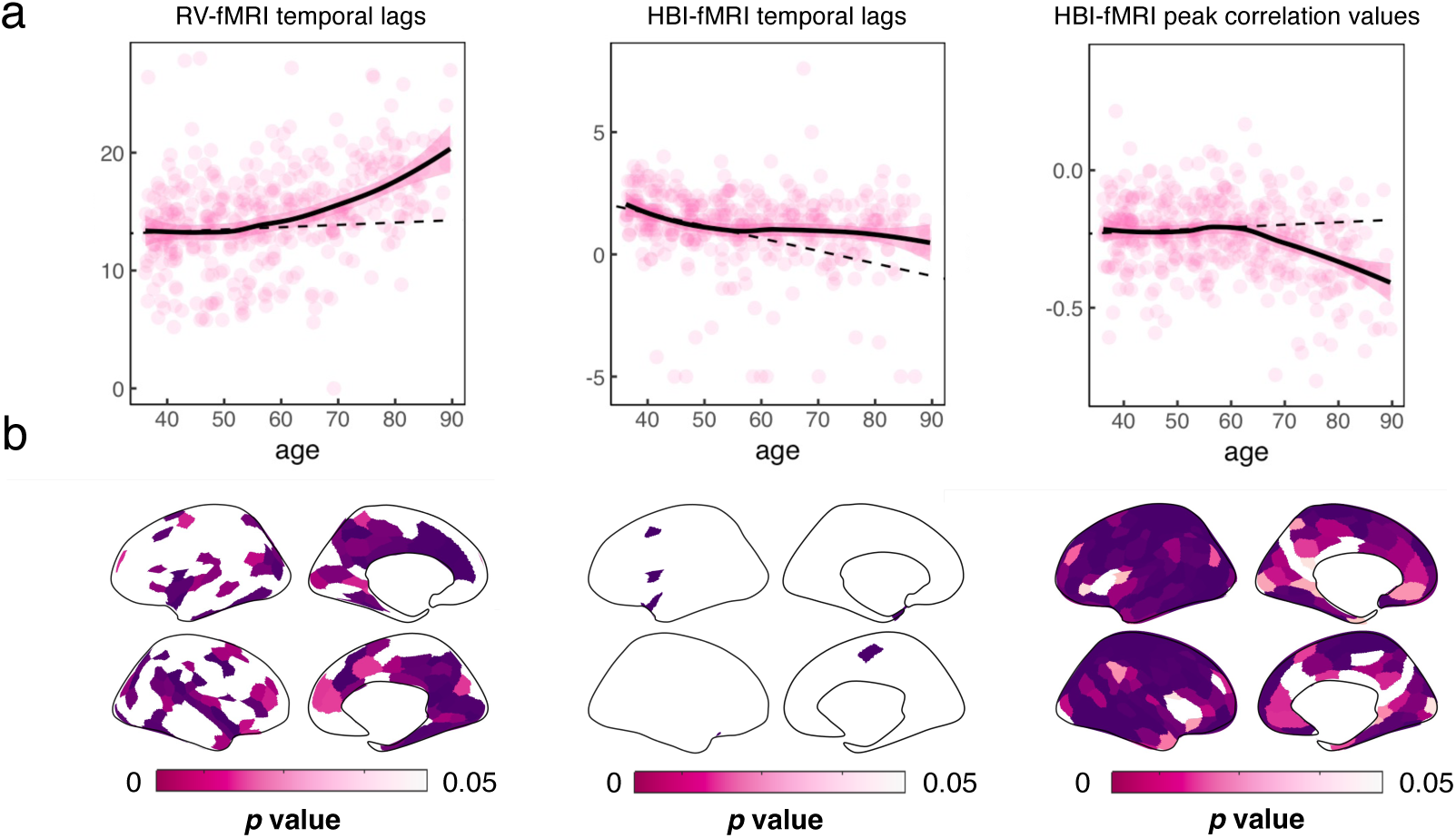
Aging rates of physiological fMRI metrics. (a) Scatterplots of RV-fMRI temporal lags, HBI-fMRI temporal lags, and HBI-fMRI peak correlation values as a function of age. Results of the global cortical gray matter are shown, and the age dependence was fitted with non-linear LOESS regression. Shaded pink regions indicate standard errors of fitting. Linear dashed lines (fitted using the Aging I group data) were included in the plots to help visualize apparent inflections in the aging rate after age sixty. (b) Regions exhibiting statistically significant changes in linear aging rates after age 60 (FDR corrected, *p* < 0.05).

When comparing the age trajectories of global physiological fMRI metrics to those of brain anatomical and perfusion features, we noticed that RV- and HBI-coupled fMRI metrics were amongst those demonstrating the most pronounced inflection in aging rates after 60 years (*Fig. 5*). This observation implies a prominent role of declined physiological functions in the later stages of brain aging and highlights the importance of considering brain-body interactions in understanding the aging process and its impact on brain health.

**Figure 5.**
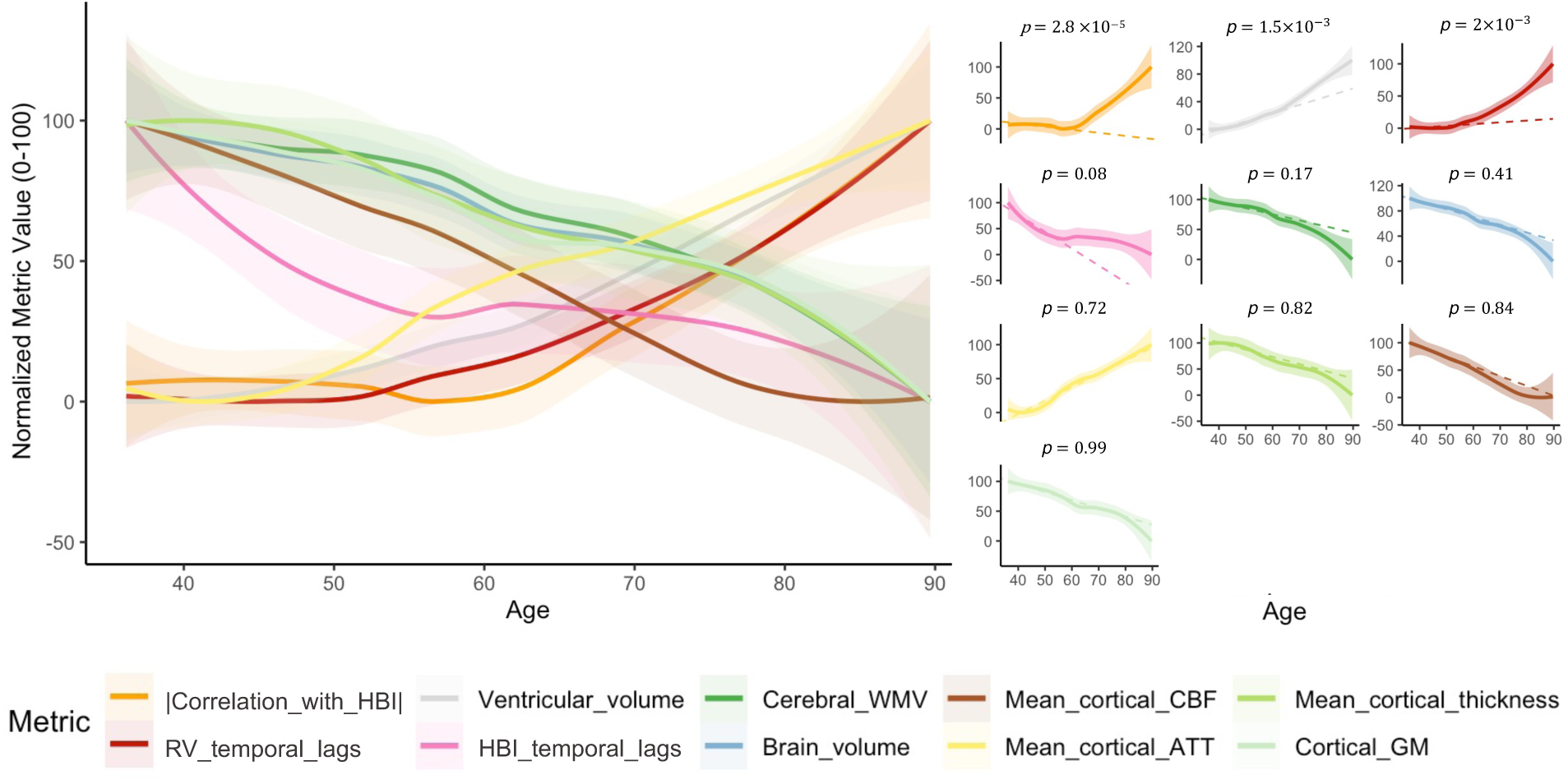
Trajectories of different physiological fMRI, structural, and tissue perfusion metrics as a function of age. Different metrics were normalized and rescaled to 0-100 for display. Results of absolute global cortical HBI-fMRI correlation values (“|Correlation_with_HBI|”), RV-fMRI temporal lags (“RV_temporal_lags”), and HBI-fMRI temporal lags (“HBI_temporal_lags”) are shown. Structural metrics include the total brain volume (“Brain_volume”), ventricular volume (“Ventricular_volume”), cerebral white-matter volume (“Cerebral_WMV”), mean cortical thickness (“Mean_cortical_thickness”), and cortical gray-matter volume (“Cortical_GM”). Brain perfusion metrics include cortical mean CBF (“Mean_cortical_CBF”) and ATT (“Mean_cortical_ATT”). For each metric, the trajectory of age dependence was fitted using LOESS, with the shaded area indicating standard errors of fitting. Individual metric fitting results are shown on the right, ranked by the statistical significance of altered linear aging rates after age 60, from the highest to the lowest. Linear dashed lines (fitted using the Aging I group data) were included in the plots to help visualize apparent inflections in the aging rate after age sixty.

## Discussion

In this study, we utilized the HCP-A resting-state dataset to characterize age-related changes in fMRI signatures of autonomic physiology. Our findings revealed significant age effects on the dynamics of physiological fMRI signals, identifying slower respiratory responses, faster cardiac responses, and stronger brain-cardiac signal coupling in aging. These physiological fMRI patterns mirror known age-related changes in vessel biomechanical properties, tissue perfusion, and autonomic function. We further demonstrated that the impact of aging on respiratory and cardiac fMRI dynamics does not progress uniformly across the lifespan, with a notable turning point around the age of 60, highlighting physiological declines in the cerebrovasculature as a key feature of aging. These results deepen our understanding of age-related changes in brain physiology and present new biomarker opportunities for investigating cardiovascular and cerebrovascular health in aging populations in future research.

Overall, the slower respiratory responses observed in our study are consistent with both prolonged arterial transit time and delayed local hemodynamic effects in aging. Aging leads to increased arteriolar tortuosity [20–22] and reduced density of the cerebral capillary and arterioles [23–27], causing both longer blood transit and arrival time, thus longer time for CO2 delivery, from lungs to the brain and from the large arteries to capillaries. The spatial consistency of age effects in ATT and RV-fMRI temporal lag patterns in gray matter (*Fig. 3b*), as well as similarly delayed responses of both measures in the white matter [30,62], all supported a prominent effect of ATT in impacting the respiratory response speed. However, it appears that ATT alone (with < 1-s lags across the cortex) cannot fully account for the differential RV-fMRI lags observed between two age groups (*Fig. 3* vs. *Fig. 1*, “Aging I vs. II”). Thus respiratory response delays may also stem partly from slower arteriolar dilation and blood delivery caused by elevations in CO2 level, i.e., slower vascular reactivity generating slower hemodynamic responses to CO2. This possibility aligns with previous evidence showing that older subjects exhibit more sluggish stimulus-driven hemodynamic responses, with a time-to-peak delay of ∼1 s compared to healthy young controls [63,64], although it should be noted that vasogenic and neurogenic responses involve distinct mechanisms of *active* vasodilation [65]. Collectively, the widespread delayed RV respiratory responses in the elder cohort may reflect a combined effect of both delayed ATTs and local vasogenic hemodynamic responses in aging. Notably, although the group-level RV-fMRI correlation values were smaller in the *Aging II* compared to the *Aging I* cohort, this between-cohort difference did not reach statistical significance (see *Supplementary Fig. S3*), which was unexpected given the reduced vascular reserve shown by previous hypercapnic studies in aging [13,31]. Since correlation values depend on the fractional contribution of the respiratory response to total fMRI variance rather than its absolute amplitude, our results suggested that total fMRI signal variance also decreases to a comparable extent with aging. This possibility was further supported by our observations that both the amplitudes of fMRI fluctuations and respiratory signals decreased in the older cohort of the HCP-A data (see *Supplementary Fig. S4*).

The accelerated global cardiac response in aging is consistent with vascular stiffening and reduced arterial compliance to the incoming cardiac pressure waves. Using phase contrast MRI [66,67] and sonography [68], previous studies have demonstrated faster pulse-wave velocity in several major vessels in old adults [69,70]. Our findings suggest that, in addition to the fast pulse wave associated with each heartbeat (∼1 Hz), slower pressure waves (<0.1 Hz) driven by heart-rate variations also propagate more rapidly with aging. These slow blood pressure waves, when transmitted to the cerebrovasculature, induce local vasodilation that displaces tissue and cerebrospinal fluid signals. Because these compartments have distinct T2* values, such displacements result in an apparent change in nominally BOLD-weighted fMRI signals. Beyond these dynamic partial volume effects, there is evidence that lower CBF could lead to faster hemodynamic responses [71,72]. Thus faster passive vasodilation may also be linked to the reduced baseline CBF seen in aging. This effect of lower baseline CBF may play a bigger role in the passive vasodilation caused by increased blood flow than in neurogenic or CO2-triggered vasogenic changes, which involve active stimulation of arteriole smooth muscle dilation that also undergoes significant alterations in aging.

The widespread increase in brain-cardiac signal coupling cannot be fully attributed to altered timing of autonomic nervous system activity in aging, in part because the associated peak timing fell outside the expected neurovascular coupling range, also because brainstem regions and cortices whose neuronal activity drives autonomic signaling (such as the insula and anterior cingulate cortex) did not exhibit the most pronounced increases in HBI-fMRI coupling (*Fig. 2c and d*). While modest contributions from in-scan arousal state differences may be present, other factors likely play a role. One possibility is that neuronal and synaptic loss, along with diminished neurovascular coupling in aging, reduces the proportion of neurogenic hemodynamic changes contributing to the total variance in fMRI signals. This reduction reflects both decreased neural activity that directly consumes oxygen [73], and impaired neurovascular coupling that *actively* controls arteriole smooth muscle dilation [74,75]. In contrast, *passive* vascular dilation in response to systemic changes in cerebral blood supply is less susceptible to deficits in neurovascular coupling. Thus this shift in neurogenic BOLD fractional contributions may elevate the correlation between fMRI and heart rate signals. Moreover, in addition to less impaired passive vasodilation, reduced cardiac outflows in aging may also be compensated by increased variability of heart rate and hypertension. Consistent with previous reports [52,53], we observed a U-shaped relationship between heart rate variability and age, with a notable increase in heart rate variability after age 60 (see *Fig. 2a* RMSSD measures and *Supplementary* Fig. 1), which has been previously attributed to heightened irregularity and fragmentation of heartbeats in aging [76] and can contribute to overall amplified brain-cardiac signal coupling. Another intertwined mechanism is hypertension common in aging [77]. Pulse pressure—the difference between systolic and diastolic pressure—showed a marked increase after age 60 in the HCP-A data (see *Supplementary* Fig. 5). This increase, together with less effective cushioning of the arterial system in aging, may increase the transmission of cardiac pressure wave energy to the cerebrovasculature [7]. This possibility is supported by a significant correlation between pulse pressure and HBI-fMRI coupling strength (see *Supplementary* Fig. 5), as well as the role of pulse pressure as a significant mediator of age-related changes in HBI-fMRI coupling (*p* = 2.4×10^-3^, tested using the *mediation* package in R).

By introducing the metric of “aging rate”, our results revealed that fMRI-based physiological metrics, particularly brain-cardiac signal coupling, do not progress linearly with age. Notably, the inflection in aging rates was more pronounced in physiological fMRI metrics compared to structural and perfusion-based imaging measures. This observation highlights altered autonomic function and vascular physiology as key markers of advanced brain aging. Furthermore, we observed spatially varying aging rates across all physiological fMRI metrics (contrasting age-related changes in *Figs. 2 and 3* with *Fig. 4*). For instance, the insula and ACC were among those exhibiting the most significant rate changes in the temporal lags of physiological responses, possibly due to their role in autonomic regulation [78] and location within cerebral arterial territories that are particularly vulnerable to late-stage aging [79].

From a methodological perspective, our study shows that resting-state fMRI signals can provide rich, multi-parametric physiological information, extending beyond the extensively studied functional connectivity estimates. These physiological fMRI metrics leverage the high spatiotemporal resolution and sensitivity of BOLD fMRI, offering new insights into certain vessel mechanical properties that are difficult to assess with conventional techniques (e.g., arterial compliance inferred from fMRI signal timing). Moreover, they eliminate the need for complex experimental setups or maneuvers required by conventional methods for assessing vascular physiology, such as gas inhalation in CVR mapping that may be impractical or unsuitable for certain patient populations. As such, physiological fMRI signals collected in the resting state hold significant clinical potential, or at least, can serve as a valuable complement to existing imaging modalities and invasive procedures for assessing brain physiology and cardiovascular health.

Finally, our study also opens up several directions that can be explored in future studies. First, while the HCP-A dataset represents a typical, healthy aging cohort, excluding common forms of atypical cognitive impairment and symptomatic stroke (see *Supplementary Fig. S6* for the cognitive test results of subjects examined in this study), it does not exclude those with common age-related vascular risks [43,44]. Therefore, the age-related changes in physiological fMRI metrics observed here may, in part, reflect the influence of prevalent vascular conditions in late-stage aging, such as atherosclerosis and hypertension discussed above. Studies with finer age range divisions (encompassing different stages of aging) and mediation analyses incorporating detailed clinical cardiovascular metrics, when available, will provide deeper insights into the complex interplay between age and vascular disease. Second, we primarily characterized physiological and biomechanical properties of the vessels by focusing on the temporal features of physiological fMRI metrics outside the neurovascular coupling range. Future investigations employing rapid fMRI sampling and multi-modal acquisitions with ground-truth neuronal information will enable further opportunities to uncover a comprehensive picture of autonomic physiology by including the neurovascular coupling components associated with focal autonomic nervous activity, such as sympathetic activity involved in regulating low-frequency heart rate variations and blood pressure waves [80–82]. Third, in this study, we examined the influence of age on respiratory and cardiac responses by separately linking RV and HBI to global and local fMRI signals, without accounting for interactions between the two processes. However, it is known that these two processes can be coupled due to mechanisms such as respiratory sinus arrhythmia or autonomic arousal [83–85], which may account for the resembling intra-cortical temporal lag patterns of RV-fMRI and HBI-fMRI correlations observed within each age group. A preliminary analysis revealed insignificant age effects in RV-HBI cross-correlations (see *Supplementary Fig. S7a*). Moreover, as detailed in *Supplementary Fig. S7b*, any potential interactions between RV and HBI are more likely to affect the early phase of respiratory fMRI responses (corresponding to the positive peak in the RV-fMRI cross-correlations in *Fig. 1a*) and the later phase of cardiac fMRI responses (corresponding to the positive peak in the HBI-fMRI cross-correlations in *Fig. 2b*), where no significant between-group differences were observed in the current dataset. Thus, we expect modest influence of RV-HBI interactions on our findings and interpretations. However, future studies might consider developing more sophisticated models that account for the intricate interactions among various physiological processes and teasing apart associated age effects respectively. Finally, sex is known to influence vascular properties through the effects of sex hormones such as estrogen and testosterone [86,87]. Significant sex differences were observed in the physiological fMRI metrics across both age groups (see *Supplementary Fig. S8*). However, at this sample size, the interaction between age and sex was not statistically significant. Future studies with larger cohorts may provide sufficient power to detect meaningful sex-specific modulation at different stages of aging, such as the impact of menopause on vessel biomechanical properties in females [88,89].

## Methods

### Data and acquisition

Functional imaging data from a subset of 400 healthy adults from the HCP-A Lifespan 2.0 Release were analyzed in this study. The selection criteria and list of subjects are detailed in *Supplementary Table S1*. Each participant underwent two eyes-open resting-state sessions (REST1 and REST2), and each session contained two ∼6-minute fMRI scans with opposite phase-encoding directions. These resting-state fMRI scans were acquired using multiband accelerated 2D echo-planar imaging (EPI): 2-mm isotropic voxels, matrix size = 104 × 104, 72 slices, no in-plane acceleration, TR = 800 ms, TE = 37 ms, flip angle = 52°, multi-band factor = 8, echo spacing = 0.58 ms. Resting-state fMRI scans from the first session (REST1_AP and REST1_PA) were analyzed in this study.

Functional imaging data from 78 healthy young adults from the HCP (3T WU-UMinn dataset) were also examined for comparison [90]. The subject IDs are listed in *Supplementary Table S2*. One 14.4-minute eyes-open resting-state BOLD fMRI scan from each subject was analyzed. The fMRI data were acquired using multiband accelerated 2D EPI: 2-mm isotropic resolution, matrix size = 104 × 90, 72 slices, no in-plane acceleration, TR = 720 ms, TE = 33.1 ms, flip angle = 52°, multi-band factor = 8, echo spacing = 0.58 ms, with the left-to-right phase encoding direction.

During these fMRI scans, concurrent cardiac signals were monitored using a fingertip pulse oximeter and respiratory signals were recorded through chest bellows, both at a sampling rate of 400 Hz.

### Extracting physiological signals from sensor recordings

An outlier-replacement filter was first applied to eliminate noise in the physiological traces due to subject motion or improper attachment of the respiratory belt or pulse oximeter. The pulse oximeter traces were then bandpass filtered (0.5–10 Hz) using a 4^th^-order Butterworth filter, and the cardiac-peak detection was performed using the MPSTD (Multi-Scale Peak and Trough Detection) detector from the open-source PPG-beats package [91].

The computation of subject-specific respiratory variation (RV) and heartbeat interval (HBI) time courses followed previous studies [3,51]. RV reflects time-dependent changes in respiratory depths and volumes; and HBI reflects the mean heartbeat intervals, i.e., the inverse of heart rates. Both metrics were computed across 6-s sliding windows centered at each measured fMRI time point.

Mean heart rate (beats per minute, BPM) and heart rate variability (measured as the root-mean-square of successive differences of normal-to-normal intervals, RMSSD [92]) were also calculated and compared between age groups.

### Preprocessing of resting-state fMRI data

All fMRI datasets underwent standard preprocessing steps, including slice-time correction and rigid-body motion realignment using AFNI [93]. Magnetic susceptibility field-induced geometric distortion was corrected using the TOPUP toolbox in FSL [94]. Quasi-periodical physiological fluctuations time-locked to cardiac and respiratory cycles were modeled using RETROICOR [95] and linearly projected out of each voxel’s fMRI time series along with six rigid-body motion parameters and low-frequency scan drifts. The initial five and the final frames of each scan dataset were removed prior to analyses. No temporal filtering was applied to the time-series data.

### Brain parcellations

All cortical and subcortical regions of interest were first delineated based on each subject’s high-resolution T1-weighted anatomical reference data. A multi-resolution functional network atlas was used to parcellate the cortical gray matter [96]. Both the 100- and 300-parcel atlases were used for fMRI data analysis. Masks of the subcortical structures, cerebellum, brainstem, and white matter were extracted from the “*aseg*” automatic segmentation generated by FreeSurfer [97]. To mitigate the potential contamination from gray-matter signals, the white-matter mask was further eroded by one voxel. All brain regions of interest (ROIs) defined in the high-resolution anatomical space were then registered to each subject’s native functional space based on the cross-modal transformation estimated by FreeSurfer’s *bbregister*.

### Cross-correlations between fMRI signals and autonomic physiology

Cross-correlations were utilized to quantify the strengths and temporal lags of fMRI responses elicited by different physiological processes. All ROI-wise fMRI signals and physiological recordings (RV and HBI) were up-sampled to a temporal resolution of 200 ms for the cross-correlation analyses, and the correlation coefficients at each time shift were Fisher-z transformed prior to group averaging. The peak/trough detection for cross-correlations was constrained to 6-s windows centered at the group mean, and a retrospective check confirmed that the vast majority of peaks/troughs fell within the set temporal boundaries. The temporal window for peak/trough detection was adjusted for white-matter and subcortical regions.

The cross-correlation analyses were performed at both global and ROI levels, and two-sample *t* tests were performed to test for significant age effects.

### Quantifying the “aging rates” of physiological metrics

To examine whether the “aging rate”, defined as the linear dependence of different physiological fMRI metrics on age, significantly differs between the two age groups, we performed the following linear regression tests. For each physiological fMRI metric, we fit it with a general linear model consisting of a constant baseline term, a binary age group indicator (“Aging I” or “Aging II” group), and an age covariate (to model the linear dependence on age). We then tested whether the inclusion of an additional conditional age covariate (i.e., derived by setting the ages of all “Aging I” participants zero) significantly improved the linear model fitting using an F test. The test results informed whether the aging rate of any specific physiological metric changed significantly after age 60.

### Relating physiological fMRI metrics to age-dependent changes in anatomical structure, perfusion, and arousal

#### Cortical thickness and gray-matter partial volume

Parcel-wise cortical thickness was quantified automatically based on the high-resolution T1-weighted anatomical reference data using FreeSurfer [97]. For each cortical parcel, the level of gray-matter volume fraction was computed by averaging across all voxels within the parcel using *mri_compute_volume_fractions* in FreeSurfer.

#### Arterial transit time and cerebral blood flow

Multi-inversion-time ASL data of the participants (*N*=400) from HCP-A were employed to compute regional ATT and CBF as detailed previously [30]. Briefly, ATT was calculated using a two-stage cross-correlation method, comparing the multi-delay ASL signal with a simulated signal based on the flow-modified Bloch equation to identify the shift that yielded the maximum correlation for each voxel. CBF was derived using a two-compartment model that factored in physiological constants and relaxation time, with results averaged over appropriate post-labeling delays.

#### Arousal effects

To track the subjects’ instantaneous arousal levels during the scans, the eye camera data from a subset of subjects (*N=88*) were also analyzed. Subjects were selected based on the visibility of the eye (minimal head coil blockage), synchronization with fMRI acquisition, and minimal interference from eyelashes or artifacts. Eye closure was detected using a customized eye detection algorithm [98], and the percentage of eye closure was calculated across 0.8-s windows (TR length) to indicate the subject’s instantaneous arousal level. The eye-closure arousal metrics were temporally shifted (*Aging I*: 6.4 s and *Aging II*: 5.6 s for all cortical parcels) and then correlated with fMRI signals to characterize arousal effects. The relative temporal lag for each group was optimized by maximizing the peak cross correlation values between the global fMRI signal and the arousal metric (see *Supplementary Fig. S2b*).

#### Spatial correlations

The spatial similarity between age-dependent changes in physiological metrics and structural, perfusion, and arousal measures were estimated using the Spearman’s correlation computed over space.

#### Trajectories of age dependence

For each physiological fMRI, structural, and perfusion metric, its trajectory across the lifespan was mapped by fitting the age dependence of each metric using local polynomial regression, implemented via the *loess* function in R.

## Acknowledgements

The authors acknowledge Amelia Strom and Richard Song for valuable discussions regarding the present results. This work was supported in part by the NIH (grants K99/R00-NS118120, U19-NS123717, U01-AG052564, U01-AG052564-S1, U54-MH091657, RF1-MH125931, and F99-AG079810), by the BrightFocus Foundation Alzheimer’s Disease Research Grant, and by the MGH/HST Athinoula A. Martinos Center for Biomedical Imaging. Computational resources were generously provided by the Massachusetts Life Sciences Center (https://www.masslifesciences.com/). Data used in the preparation of this article were obtained from the National Institute of Mental Health (NIMH) Data Archive (NDA). NDA is a collaborative informatics system created by the National Institutes of Health to provide a national resource to support and accelerate research in mental health. Dataset identifier(s): 10.15154/f1ma-9x37. This manuscript reflects the views of the authors and may not reflect the opinions or views of the NIH or of the Submitters submitting original data to NDA.

## Supplementary Materials

**Table S1:**
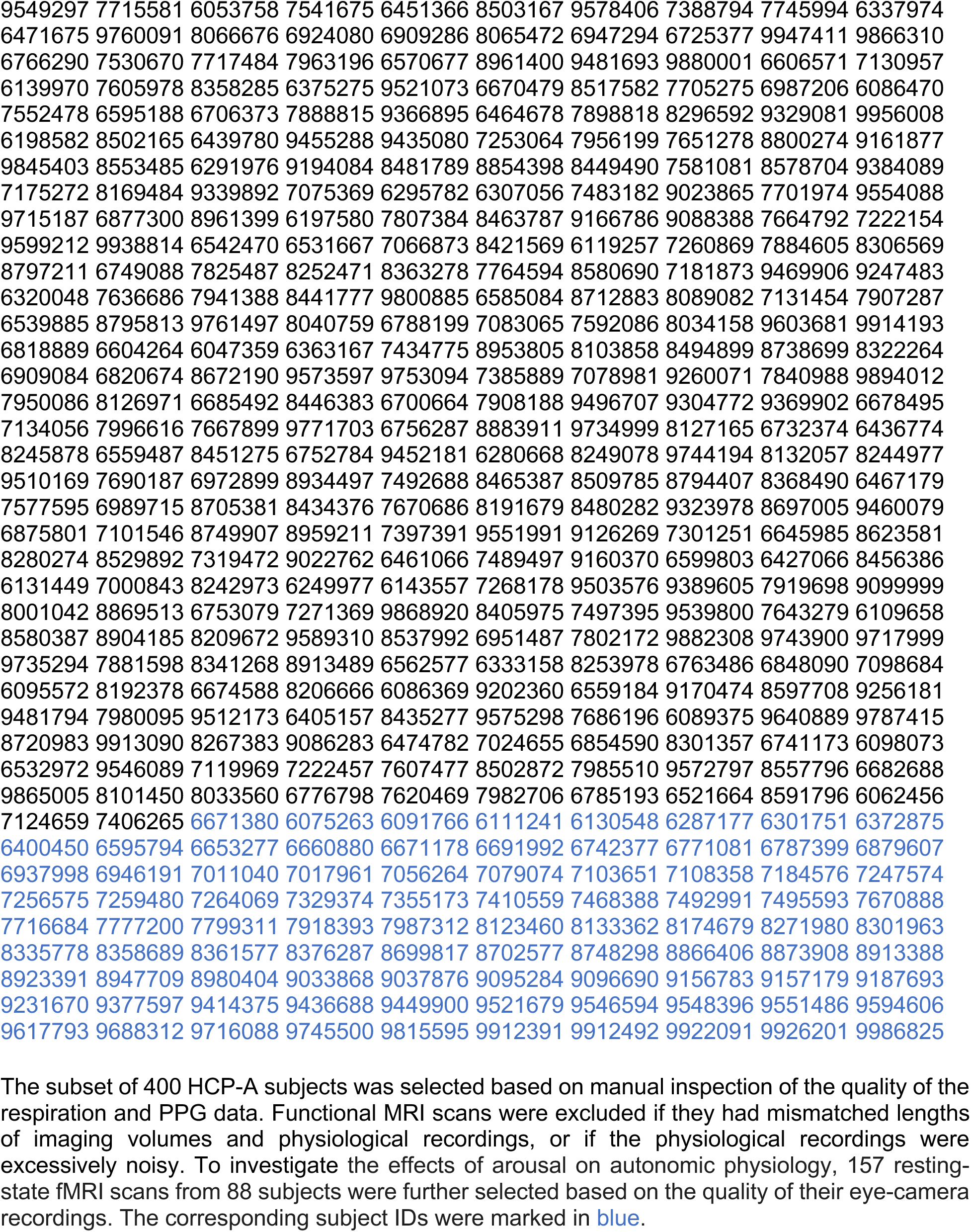
ID numbers of the HCP-A subjects included in the present study:

**Table S2:**
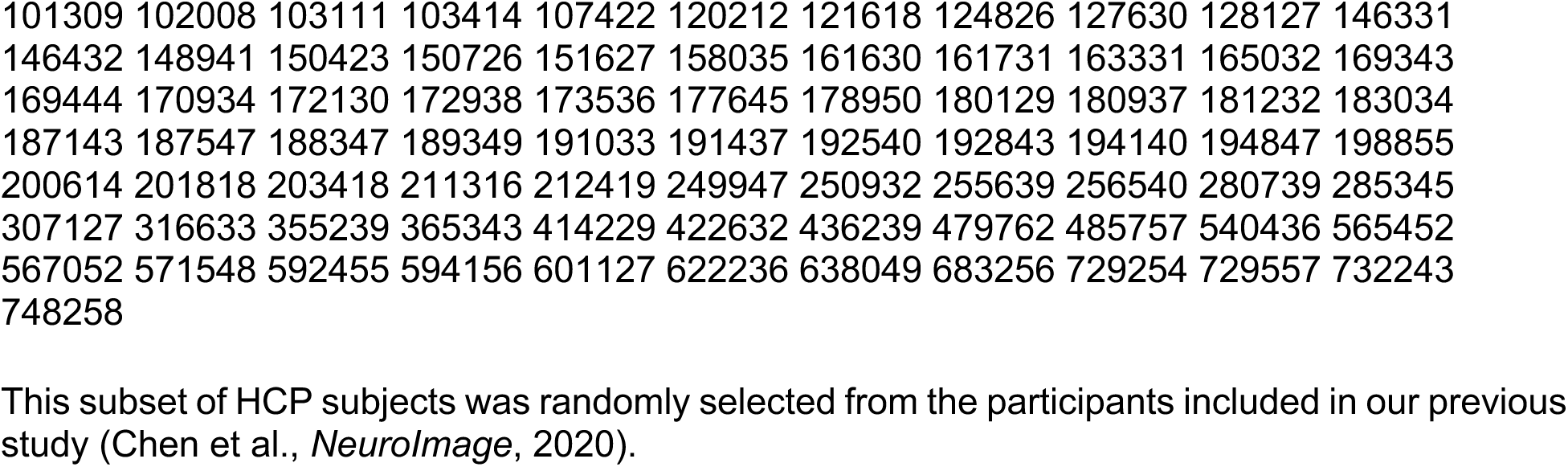
ID numbers of the HCP young healthy subjects included in the present study:

**Figure S1:**
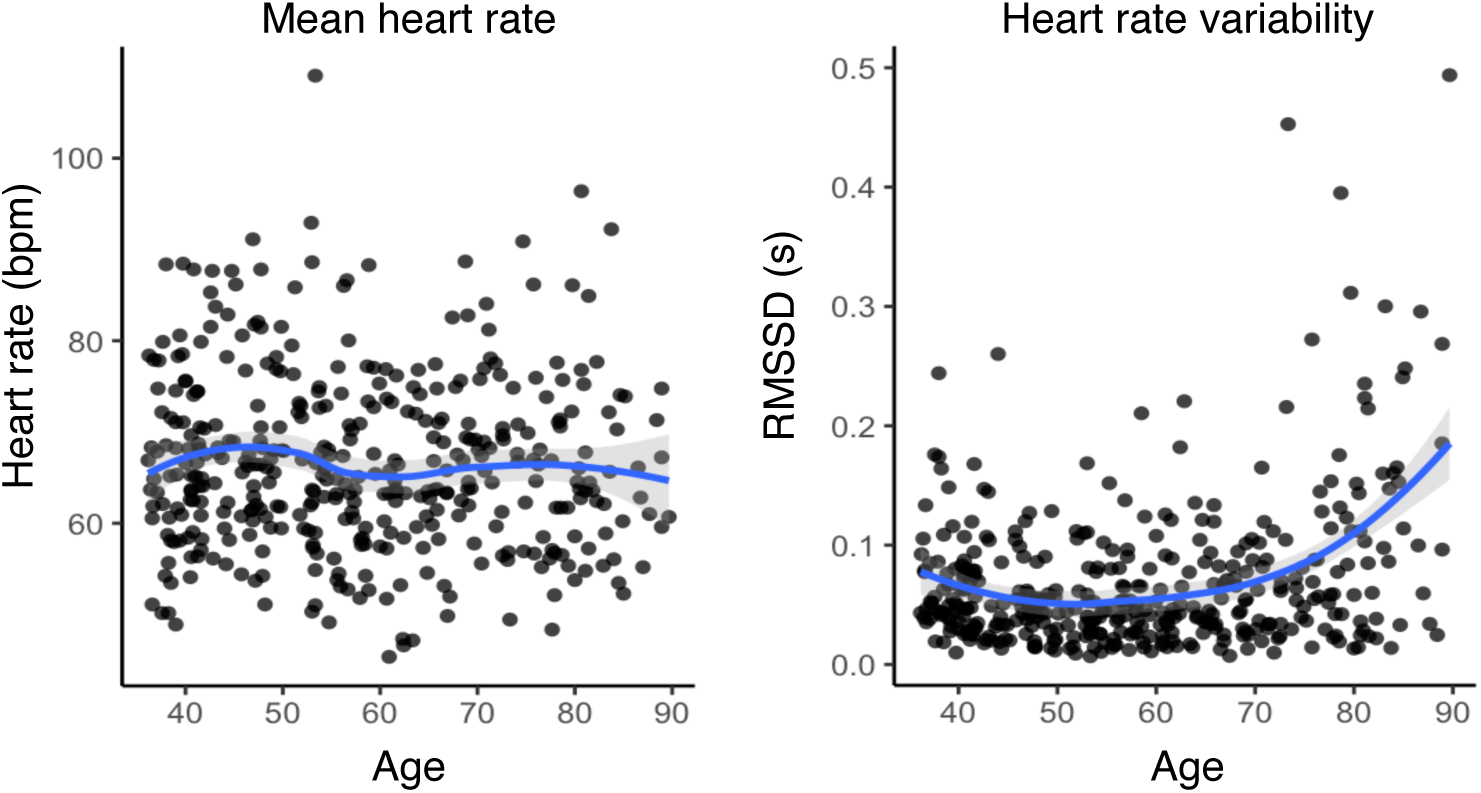
Mean heart rate and heart rate variability as a function of age. Each dot represents the results of one subject; the trajectories of mean heart rate and heart rate variability across the lifespan were mapped by fitting the age dependence using local polynomial regression, implemented via the *loess* function in R, with the shaded area indicating standard errors of fitting.

**Figure S2:**
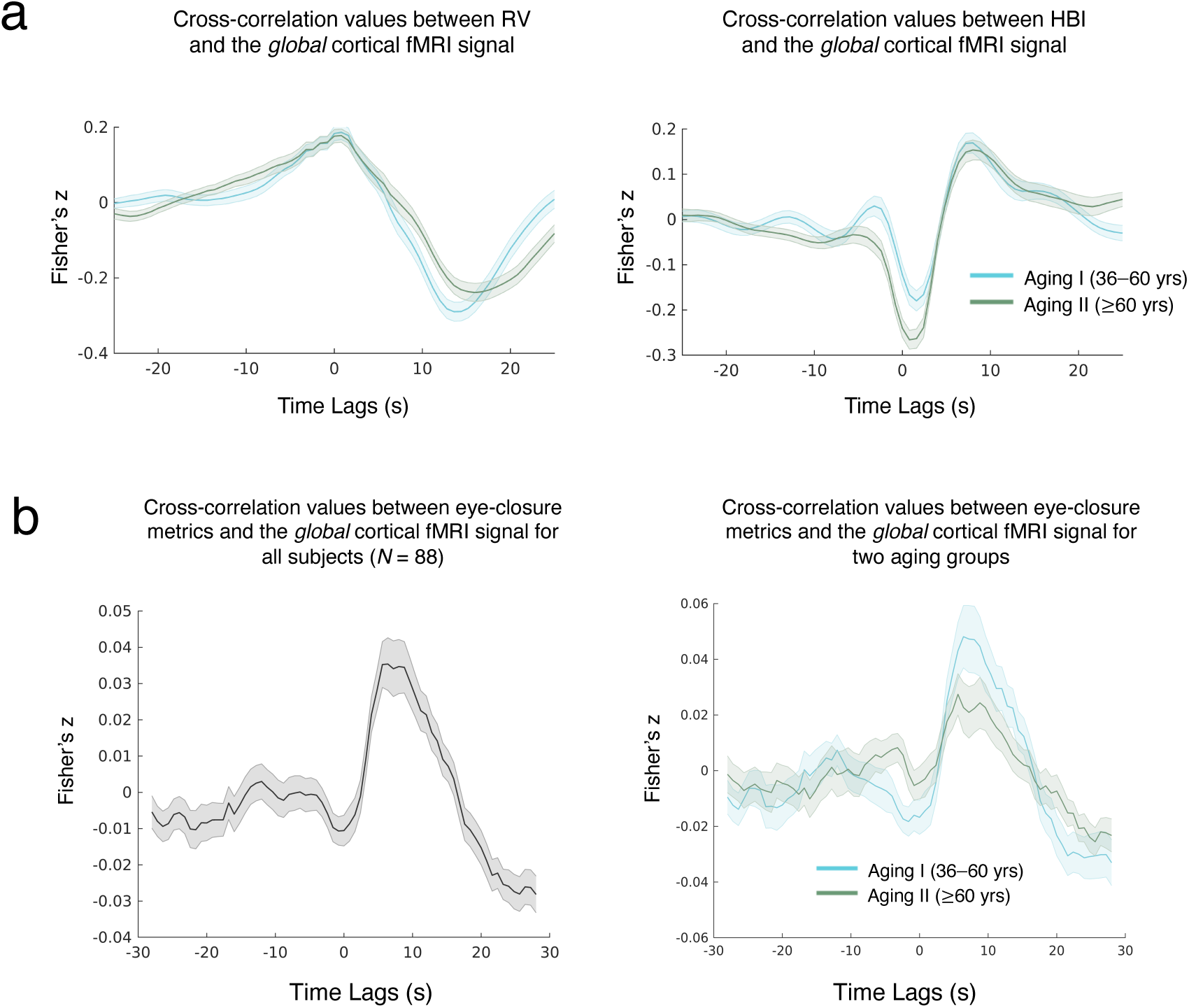
Cross-correlations between fMRI signals and RV/HBI and eye-closure arousal (*N*=88). (a) Replication of the RV/HBI-fMRI cross-correlation results in Figs. 1a & 2b, using the subset of data for which in-scan arousal level was assessed based on eye-camera data; (b) Cross-correlations between global fMRI signals and eye-closure metrics.

**Figure S3:**
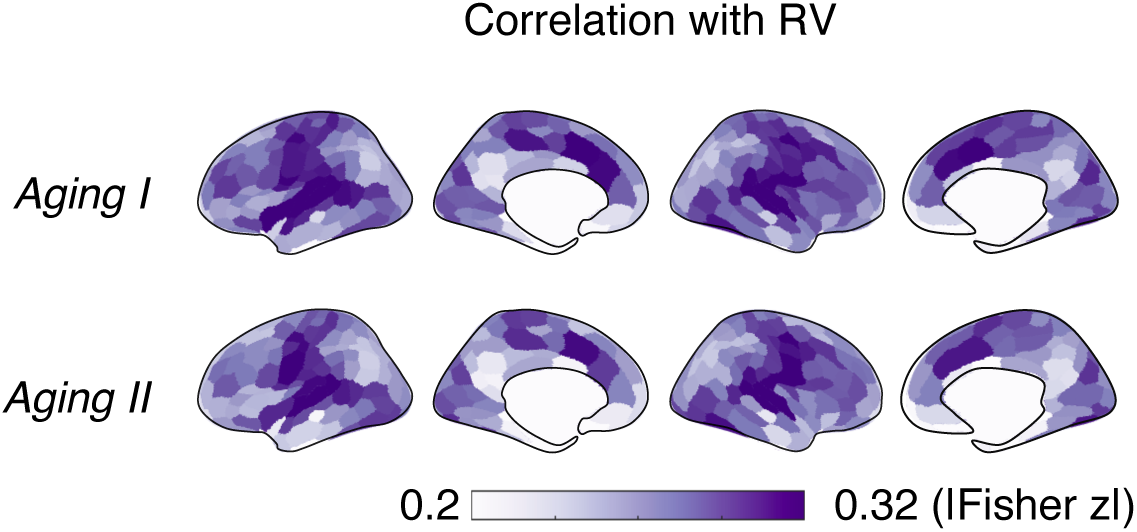
Age-related changes in the strengths of RV-fMRI coupling. Spatial distributions of region-specific couplings between RV and fMRI signals (based on the Schaefer 300-parcel atlas and the trough of the temporal cross-correlations, Fig. 1b). For both age groups, somatosensory motor, visual and dorsal attention region exhibited the strongest RV-fMRI coupling. Yet, no regions showed statistically significant between-group differences (“Aging II > I”, *p* < 0.05, FDR).

**Figure S4:**
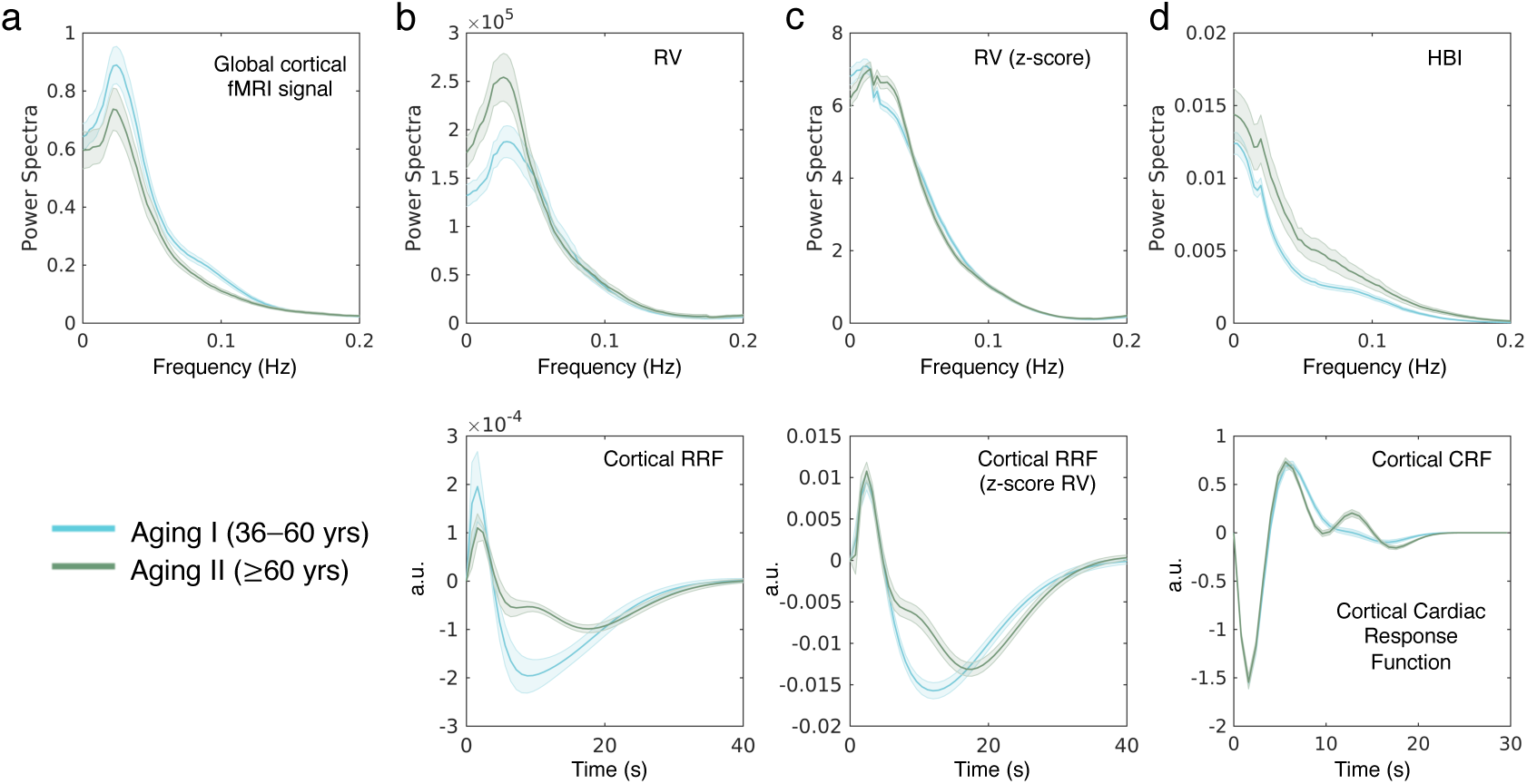
Age-related changes in the power spectra of fMRI signals, RV, HBI, and physiological fMRI responses. (a) Power spectra of percent signal changes of the global cortical fMRI signal. (b) Power spectra of RVs and the mean cortical respiratory response functions (RRFs). (c) Power spectra of RVs and the mean cortical RRFs, estimated after converting the individual RV to z-scores. This additional z-score normalization was performed because RV is sensitive to the placement of the respiratory below and less quantitative. Age-related reductions in RRFs persisted, albeit became smaller, after the additional normalization step. (d) Power spectra of HBIs and the mean cortical cardiac response function (CRFs). RRFs and CRFs were derived by fitting two sets of basis functions (five basis functions for each physiological response) to fMRI data using ordinary least squares fitting, as detailed in Chen et al., *NeuroImage*, 2020.

**Figure S5:**
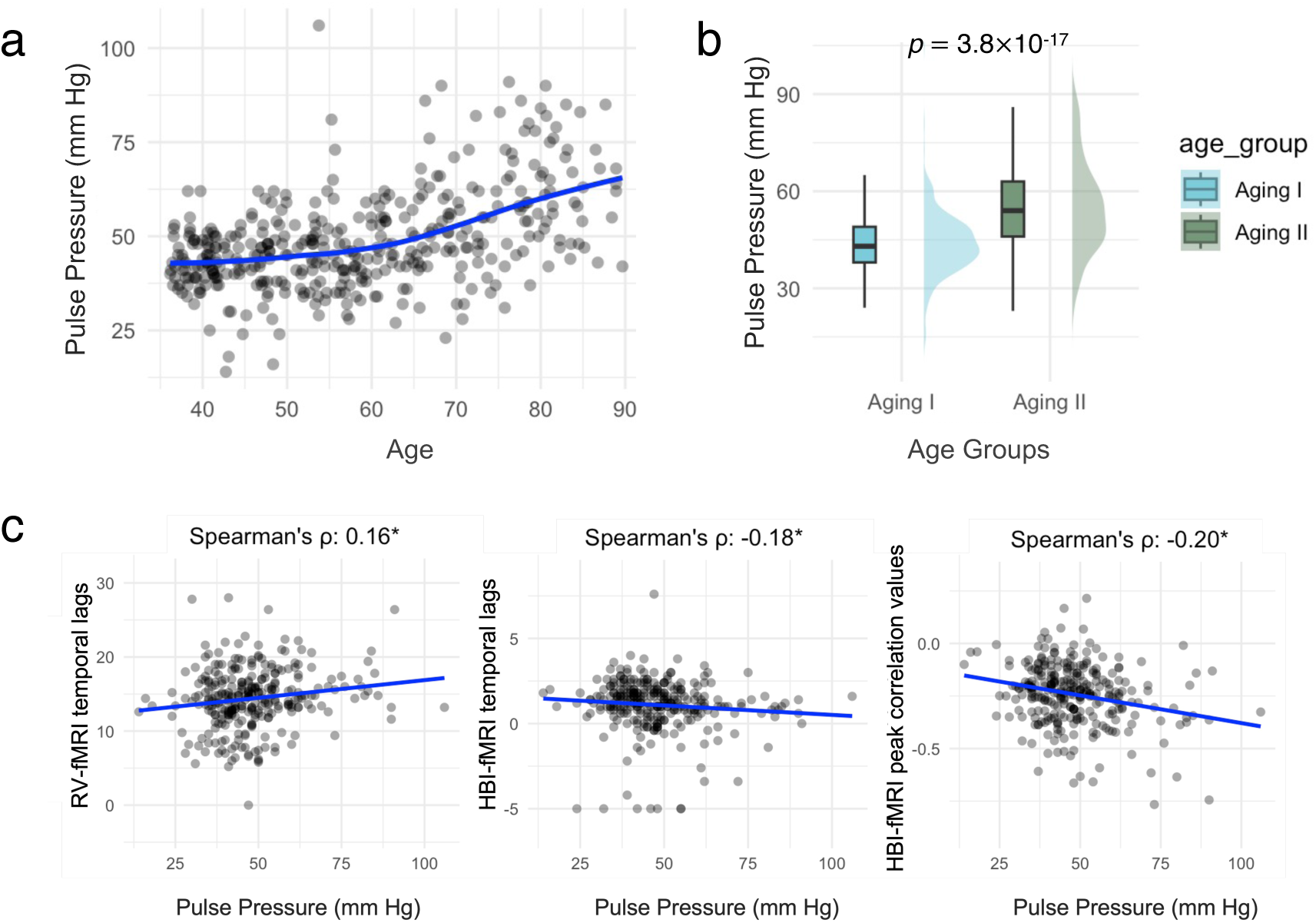
Age-related changes in pulse pressure, and its association with physiological fMRI metrics. (a) Pulse pressure (calculated as the difference between systolic and diastolic blood pressures) as a function of age, each dot represents the results of a single subject. (b) Mean and standard deviations of pulse pressures within each age group. (c) Spearman’s correlation between pulse pressure and different physiological fMRI metrics. * denotes *p* < 0.05.

**Figure S6:**
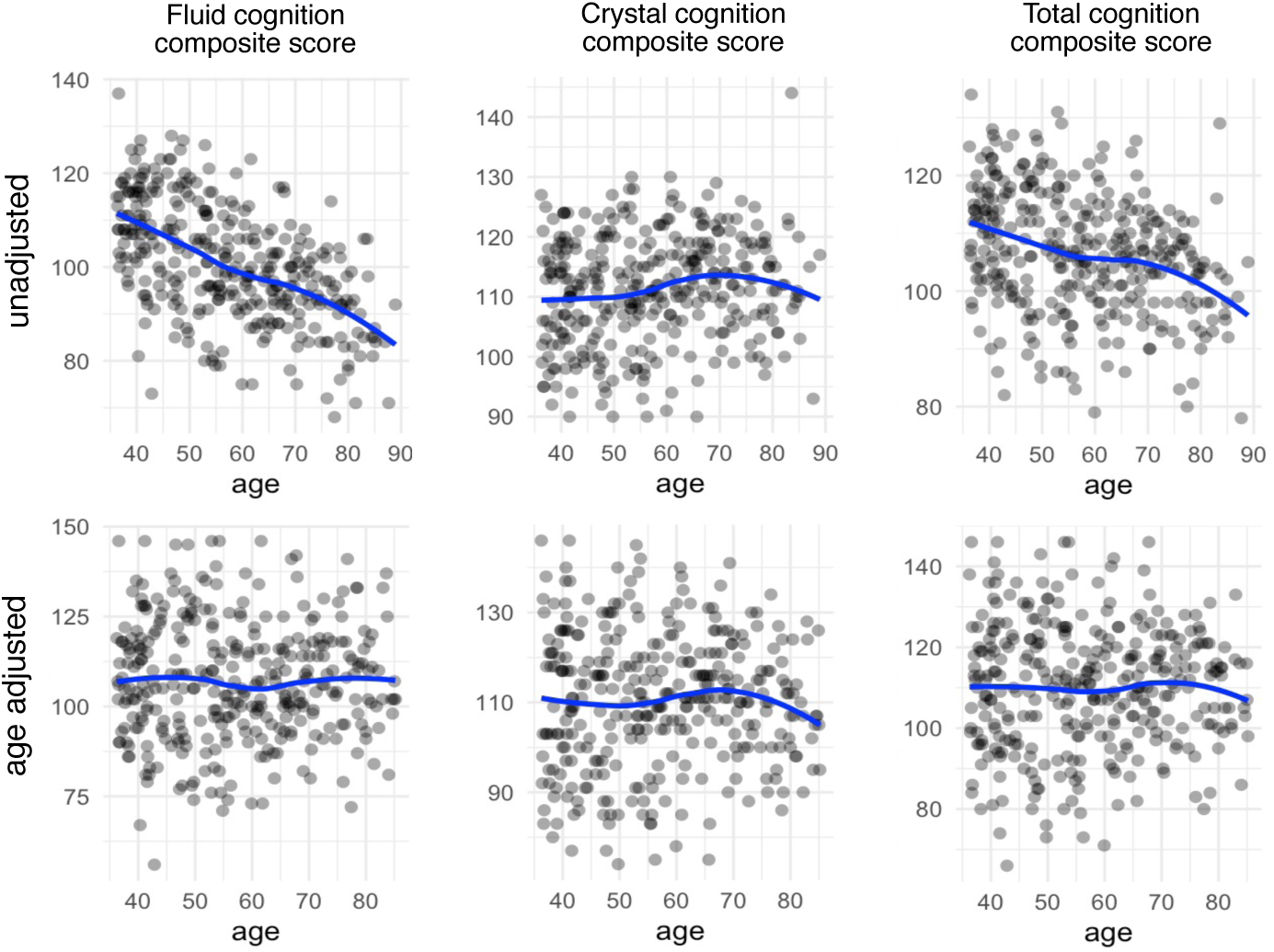
Age-related changes in cognitive scores. As expected, subjects’ cognitive scores decline with age; however, after adjusting for age, they remain within the typical range, indicating low cognitive impairments in HCP-A subjects.

**Figure S7:**
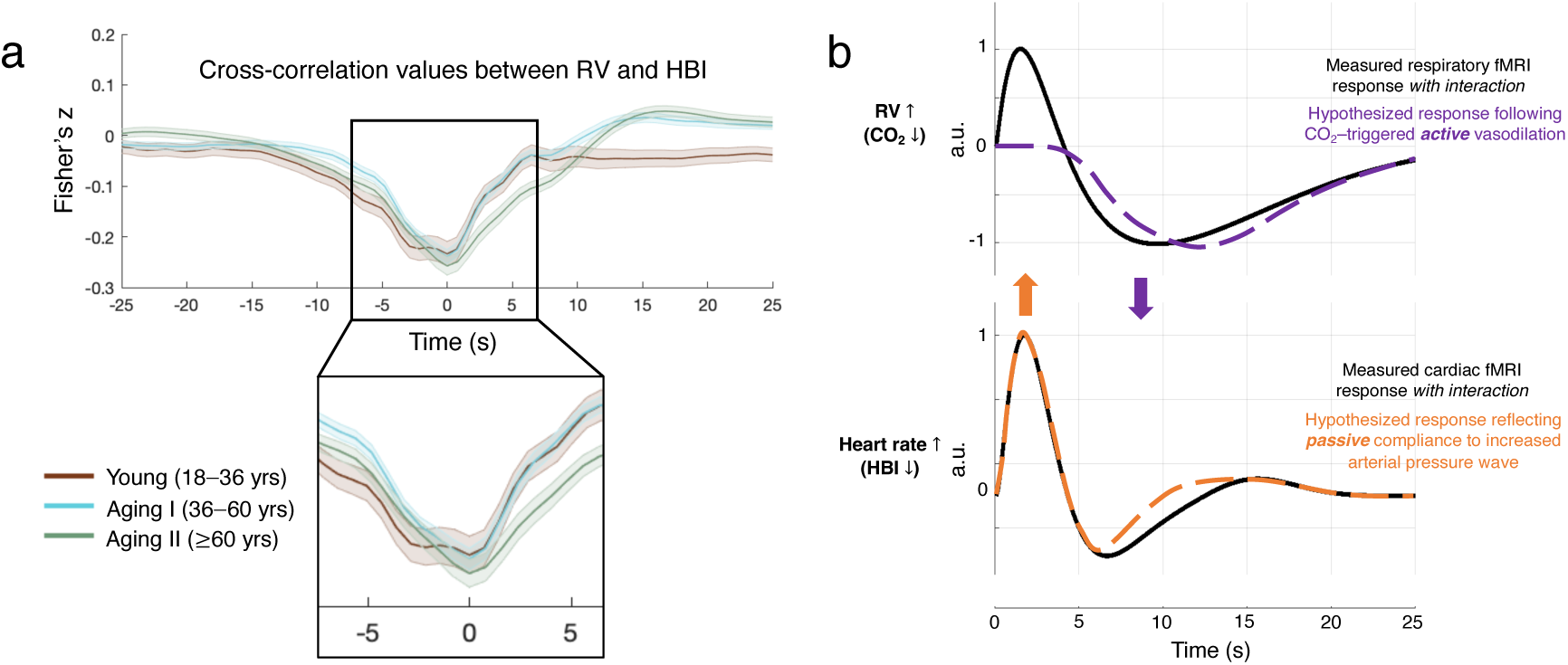
Influence of RV-HBI coupling on physiological fMRI responses. (a) Fisher’s z-transformed cross-correlation between RV and HBI at varying time lags (seconds) for three distinct age groups. The shaded areas indicate the standard errors across subjects within each age group. An inset highlights the central region (-5 to 5 seconds) to better visualize the trough correlation values of three age groups. (b) Hypothesized influence of RV-HBI interactions on measured respiratory and cardiac fMRI responses, illustrated using cortical physiological fMRI responses from Chen et al., *NeuroImage*, 2020. Arrows indicate possible interactions: fMRI responses driven by increased arterial pressure waves (orange) may contribute to the early peaks in measured respiratory fMRI responses; while the trough of CO2 responses (purple) may influence the later phase of the measured cardiac fMRI responses.

**Figure S8:**
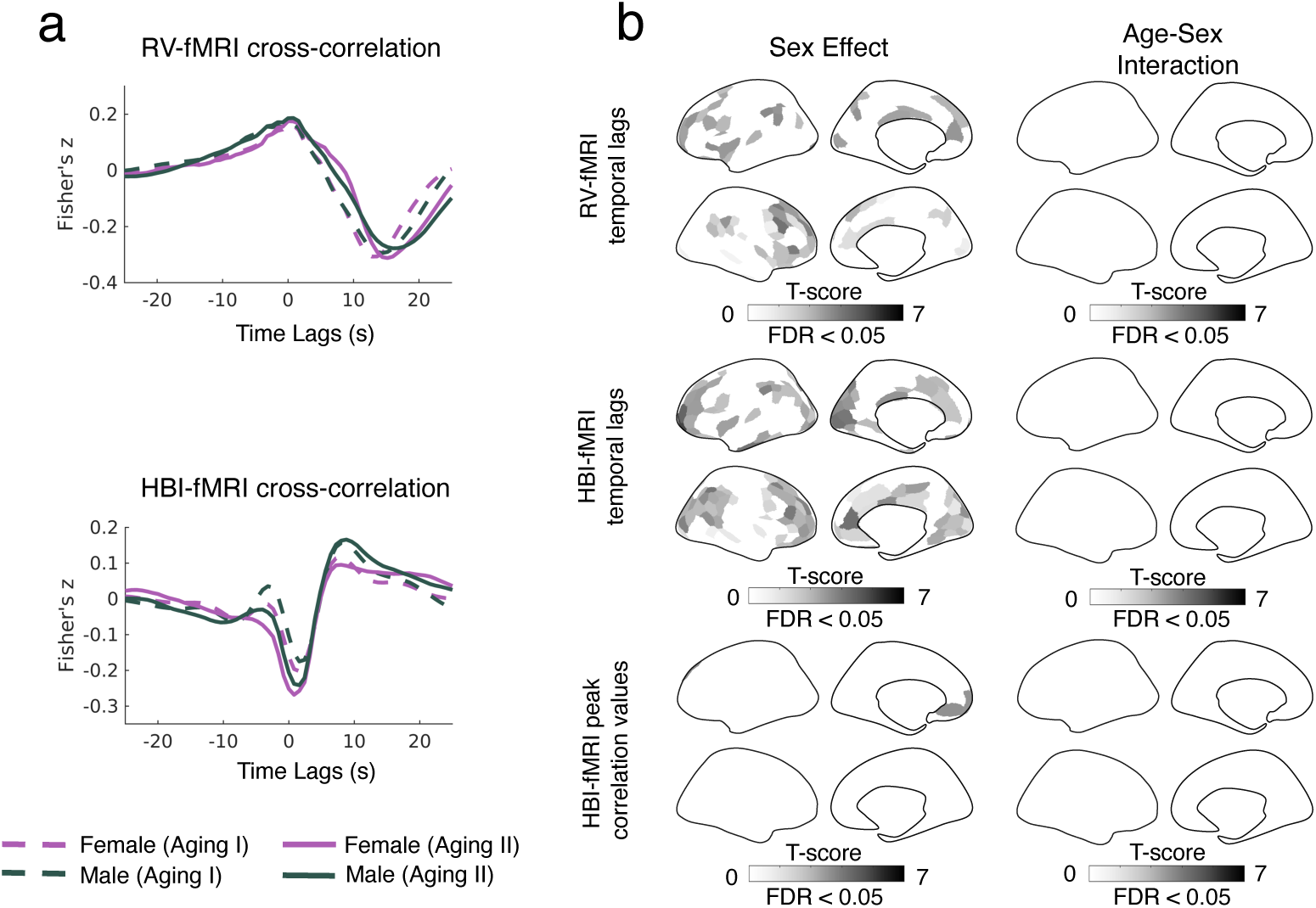
Sex effects in the physiological fMRI metrics. (a) RV/HBI-fMRI cross-correlation results, averaged across subjects within each sex and age group. (b) “Sex Effect”: spatial distributions of cortical regions showing significant sex differences; “Age-Sex Interaction”: no regions exhibited significant age-sex interactions for any of the three physiological fMRI metrics.

## Notes

### Competing Interest Statement

The authors have declared no competing interest.

